# Inter-Species Differences in the Response of Sinus Node Cellular Pacemaking to Changes of Extracellular Calcium

**DOI:** 10.1101/771972

**Authors:** Axel Loewe, Yannick Lutz, Norbert Nagy, Alan Fabbri, Christoph Schweda, András Varró, Stefano Severi

## Abstract

Changes of serum and extracellular ion concentrations occur regularly in patients with chronic kidney disease (CKD). Recently, hypocalcemia, i.e. a decrease of the extra-cellular calcium concentration [*Ca*^2+^]_*o*_, has been suggested as potential pathomechanism contributing to the unexplained high rate of sudden cardiac death (SCD) in CKD patients. In particular, there is a hypothesis that hypocalcaemia could slow down natural pacemaking in the human sinus node to fatal degrees. Here, we address the question whether there are inter-species differences in the response of cellular sinus node pacemaking to changes of [*Ca*^2+^]_*o*_. Towards this end, we employ computational models of mouse, rabbit and human sinus node cells. The Fabbri et al. human model was updated to consider changes of intracellular ion concentrations. We identified crucial inter-species differences in the response of cellular pacemaking in the sinus node to changes of [*Ca*^2+^]_*o*_ with little changes of cycle length in mouse and rabbit models (<83 ms) in contrast to a pronounced bradycardic effect in the human model (up to > 1000 ms). Our results suggest that experiments with human sinus node cells are required to investigate the potential mechanism of hypocalcaemia-induced bradycardic SCD in CKD patients and small animal models are not well suited.

## I. INTRODUCTION

The extracellular milieu in mammals is tightly controlled, i.e. amongst others the concentration of the essential electrophysiological ions is kept almost constant under physiological conditions. In disease conditions such as chronic kidney disease (CKD), the body can fail to regulate the ion concentrations including [*K*^+^]_*o*_ and [*Ca*^2+^]_*o*_. Patients suffering from CKD have an exceptionally high risk of sudden cardiac death (SCD). Indeed, it is 14x higher than in cardiovascular patients with normal kidney function [1]. This high rate in CKD patients cannot be explained by traditional SCD risk factors and recent clinical studies using implantable loop recorders point towards severe bradycardia and asystole as the prevailing arrhythmias causing SCD in CKD patients [2], [3]. An initial computational study from our group suggests a slowing of the cellular pacemaking rate in the human sinus node caused by reduced [*Ca*^2+^]_*o*_ as a likely mechanism causing bradycardic SCD in CKD patients [4]. In vitro confirmation of this in silico finding is required. However, it is not clear which experimental setting (i.e. human vs. animal sinus node cells) is suitable considering the marked difference in pacemaking rate (mouse: 500bpm, rabbit: 300bpm, dog: 100bpm; human: 60bpm). Therefore, we study inter-species differences of sinus node cellular pacemaking response to [*Ca*^2+^]_*o*_ changes using computational models of rabbit, mouse, and human sinus node cells together with rabbit experimental data.

## II. METHODS

### A. Model Extension

The original Fabbri et al. model [5] considers constant intracellular *K*^+^ and *Na*^+^ concentrations, which is a valid assumption during cyclic steady-state simulations at the reference extracellular concentrations. However, the intracellular concentrations are likely to respond to changes of the extracellular concentrations. Therefore, we extended the model to account for the dynamic intracellular *K*^+^ and *Na*^+^ balance based on the influx and efflux of ions carrying the transmembrane currents. Moreover, we included a current through the small conductance calcium-activated potassium channel (*I_SK_*) as proposed before [6], [7]. Lastly, we considered the dependance of the maximal *I_Kr_* conductance *g_Kr_* on [*K*^+^]_*o*_ as described before for other cardiomyocytes [8]:

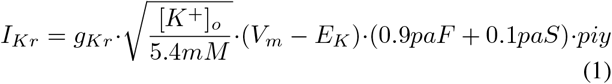

After introducing the changes to the model equations, the parameters in the upper part of Table I were optimized with respect to a cost function considering the experimental values in Table II and the existence of a cyclic steady-state similar to the method used for the original Fabbri et al. model detailed in [5]. Table I lists the optimized parameter values, Table III gives the initial values. The updated model was validated against experimental action potential (AP) and calcium transient (CaT) recordings as well as the effect of mutations similarly to the validation of the original model [5].

**TABLE I.** SUGGESTED PARAMTER CHANGES FOR THE FABBRI ET AL. HUMAN SINUS NODE CELL MODEL [5].

| Parameter | Original model | Updated model | Change |
| --- | --- | --- | --- |
| i_NaK_max (nA) | 8.11e-02 | 1.37e-01 | +68.9% |
| P_CaL (nA/mM) | 4.58e-01 | 5.05e-01 | +10.3% |
| k_dl (mV) | 4.34 | 5.85 | +34.8% |
| V_dl (mV) | -16.45 | -7.77 | -52.8% |
| k_fl (mV) | 5.30 | 5.17 | -2.5% |
| V_fl (mV) | -37.40 | -12.18 | -67.4% |
| g_Kur (μS) | 1.54e-04 | 7.06e-05 | -54.2% |
| g_to (μS) | 3.50e-03 | 1.67e-03 | -52.3% |
| g_Kr (μS) | 4.24e-03 | 4.99e-03 | +17.7% |
| g_Ks (μS) | 6.50e-04 | 8.64e-04 | +32.9% |
| g_fK (μS) | 2.68e-03 | 1.17e-02 | +337% |
| g_fNa (μS) | 1.59e-03 | 6.96e-03 | +337% |
| EC50_SK (mM) | - | 7.00e-04 | new |
| n_SK | - | 2.2 | new |
| g_SK (μS) | - | 1.65e-04 | new |

The small conductance calcium-activated potassium (SK) channel was newly introduced. Moreover, the model was extended by considering $g_{Kr}$ dependance on $[K^+]_o$ and dynamic changes of $[Na^+]_i$ and $[K^+]_i$ .

**TABLE II.** COMPARISON OF ACTION POTIENTAL (AP) AND CALCIUM TRANSIENT (CAT) FEATURES.

| Marker | Experimental [13] | Original model [5] | Updated model |
| --- | --- | --- | --- |
| MDP (mV) | -61.7 ± 4.3 | -58.9 | -60.9 |
| cycle length (ms) | 828 ± 15 | 815 | 828 |
| $(dV/dt)_{max}$ (V/s) | 4.6 ± 1.2 | 7.3 | 6.5 |
| APD<sub>20</sub> (ms) | 64.9 ± 16.9 | 86 | 73.8 |
| APD<sub>50</sub> (ms) | 101.5 ± 27.0 | 124 | 108.2 |
| APD<sub>90</sub> (ms) | 143.5 ± 34.9 | 149.3 | 130.4 |
| AP overshoot (mV) | 16.4 ± 0.7 | 26.4 | 26.8 |
| DDR<sub>100</sub> (mV/s) | 48.9 ± 18 | 48.1 | 75.7 |
| $[Ca]_{i,min}$ (nM) | 105 | 84 | 111 |
| $[Ca]_{i,max}$ (nM) | 220 | 189 | 263 |
| CaT amplitude (nM) | 115 | 105 | 152 |
| CaTD<sub>20</sub> (ms) | 138.9 | 136 | 117 |
| CaTD<sub>50</sub> (ms) | 217.4 | 206 | 165 |
| CaTD<sub>90</sub> (ms) | 394.0 | 552 | 478 |

Abbreviations: maximum diastolic potential (MDP); AP duration (APD); diastolic depolarization rate over the first 100 ms of diastolic depolarization (DDR<sub>100</sub>); CaT duration (CaTD).

**TABLE III.** INITIAL VALUES OF THE 33 ORDINARY DIFFERENTIAL EQUATIONS CONSIDERED IN THE UPDATED HUMAN SINUS NODE CELL MODEL.

| Parameter | Initial value | Parameter | Initial value |
| --- | --- | --- | --- |
| $V_m$ (mV) | -46.19 | $[Ca]_{nsr}$ (mM) | 6.69e-01 |
| $[Ca]_{sub}$ (mM) | 8.60e-05 | $[Ca]_i$ (mM) | 1.27e-04 |
| $Na_i$ (mM) | 6.08 | $fTMM$ | 5.92e-01 |
| $K_i$ (mM) | 139.14 | $fCMi$ | 2.78e-01 |
| $y$ | 1.03e-02 | $fCMs$ | 2.07e-01 |
| $m$ | 3.78e-01 | $fTC$ | 2.47e-02 |
| $h$ | 5.20e-03 | $fTMC$ | 3.30e-01 |
| $dL$ | 1.40e-03 | $fCQ$ | 1.88e-01 |
| $fL$ | 9.95e-01 | $rKur$ | 9.06e-03 |
| $fCa$ | 7.82e-01 | $s$ | 8.66e-01 |
| $dT$ | 1.92e-01 | $q$ | 4.74e-01 |
| $fT$ | 3.75e-02 | $r_{to}$ | 1.25e-02 |
| $RCa_{rel}$ | 9.00e-01 | $paS$ | 3.45e-01 |
| $OCa_{rel}$ | 1.75e-08 | $paF$ | 8.34e-04 |
| $ICa_{rel}$ | 1.88e-09 | $piy$ | 7.37e-01 |
| $RICa_{rel}$ | 9.68e-02 | $n$ | 1.06e-01 |
| $[Ca]_{jsr}$ (mM) | 5.87e-01 | | |

### B. Inter-Species Comparison

To compare the effect of [*Ca*^2+^]_*o*_ changes on sinus node cellular pacemaking between species, we used the Garny et al. [9] and Severi et al. [10] rabbit models as well as the Kharche et al. mouse model [11] besides the original and updated human models described above. In all of the models, [*Ca*^2+^]_*o*_ was varied between 0.9 and 2.9 mM. The coupled systems of ordinary differential equations were numerically integrated for 100 s to reach a cyclic steady-state.

As wetlab data on this phenomenon are very sparse, we performed experiments in isolated sinus node tissue from young New-Zealand white rabbits at 37°C to complement our computational analyses. The protocol was approved by the Review Board of the Department of Animal Health and Food Control of the Ministry of Agriculture and Rural Development, Hungary (XIII./1211/2012). The details of the conventional microelectrode technique were described before [12]. The calcium content of the bath solution was varied (0.9 / 1.35 / 1.8 mM) and the cycle length of spontaneous beating rate was determined from AP recordings in n=6 rabbits.

## III. RESULTS

### A. Updated Model

Table II compares AP and CaT features of the updated model to the original model as well as to the experimental data by Verkerk et al. [13]. AP features of the updated model were closer to experimental values than those of the original model except for APD_90_, AP overshoot, and the diastolic depolarization rate during the first 100 ms (DDR_100_) which is higher in the updated model leading to a more pronounced biphasic diastolic deploarization. The CaT was of higher amplitude and shorter duration at 20% and 50% repolarization but longer duration at 90% repolarization. The effect of *I_f_*, *I_Na_*, and *I_Ks_* mutations (Fig. 1) was not markedly affected by the changes to the model with a tendency towards bigger effects for HCN4 mutations and smaller effects for SCN5A mutations. The response to complete *I_f_* block (cycle length +25.9%) was in accordance with the available experimental human data (+26% [13]).

**Fig. 1.**
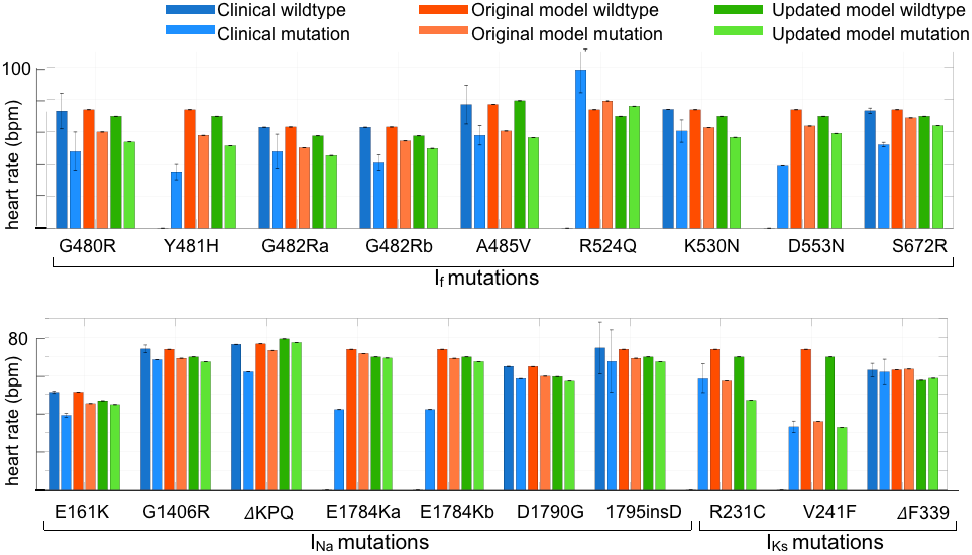
Comparison of clinically observed and simulated effects of HCN4 (*I_f_*), SCN5A (*I_Na_*), and KCNQ1 (*I_Ks_*) mutations. Details regarding the clinical data and implementation of the mutations in the human sinus node cell model were presented with the original model [5]. No clinical data from non-carrier family members were available for the Y481H, R524Q, D553N *I_f_* mutations and E1784K *I_Na_* mutations. Standard deviation of clinical data are indicated where available. Clinical data from [5].

### B. Inter-Species Comparison

Fig. 2A shows how the different models respond to changes of [*Ca*^2+^]_*o*_. While there is a consistent trend towards longer cycle length, i.e. bradycardia, under hypocalcaemic conditions, it is intriguingly stronger in the human sinus node cell models (Fig. 2A, Fig. 3A). In contrast to the original Fabbri et al. model, the extensions introduced in this work attenuate the bradycardic effect of hypocalcaemia indicating that variable intracellular ion concentrations can counterbalance the effect of decreased [*Ca*^2+^]_*o*_ to some extent, although with a marked remaining effect of −40.7 bpm when reducing [*Ca*^2+^]_*o*_ from 1.8 to 0.9 mM. Pacemaking brakes down for [*Ca*^2+^]_*o*_ < 1.2 mM in the original Fabbri et al. model. The rabbit experimental data match the rabbit model results (Fig. 3A). Decreasing [*Ca*^2+^]_*o*_ causes higher (absolute) maximum diastolic potential (MDP) consistently across species with the exception of the Garny et al. rabbit model, which is insensitive (Fig. 2B). Decreasing [*Ca*^2+^]_*o*_ has little effect on DDR_100_ in the rabbit models whereas spontaneous depolarization becomes slower in the human models and, most pronounced, in the Kharche et al. mouse model (Fig. 2C). APD_90_ is insensitive to [*Ca*^2+^]_*o*_ changes in the mouse model and showed an inverse relation in the Garny et al. rabbit model. In contrast, the Severi et al. rabbit model and the human models exhibit a direct relation (Fig. 2D, Fig. 3C).

**Fig. 2.**
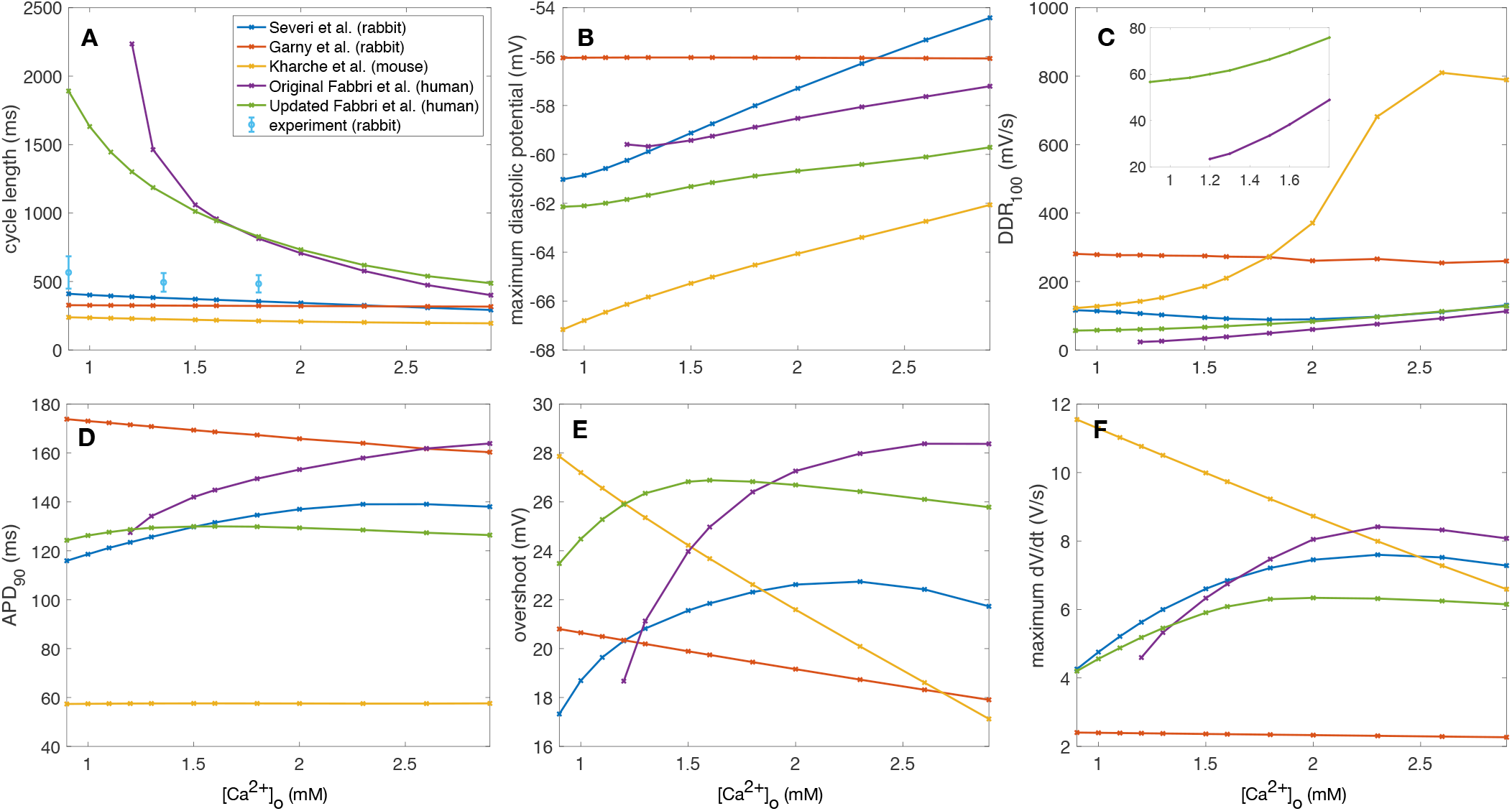
Effect of extracellular calcium concentration variation on action potential properties in sinus node cellular models of different species. Abbreviations: diastolic depolarization rate over the first 100 ms of diastolic depolarization (DDR_100_), action potential duration at 90% repolarization (APD_90_).

**Fig. 3.**
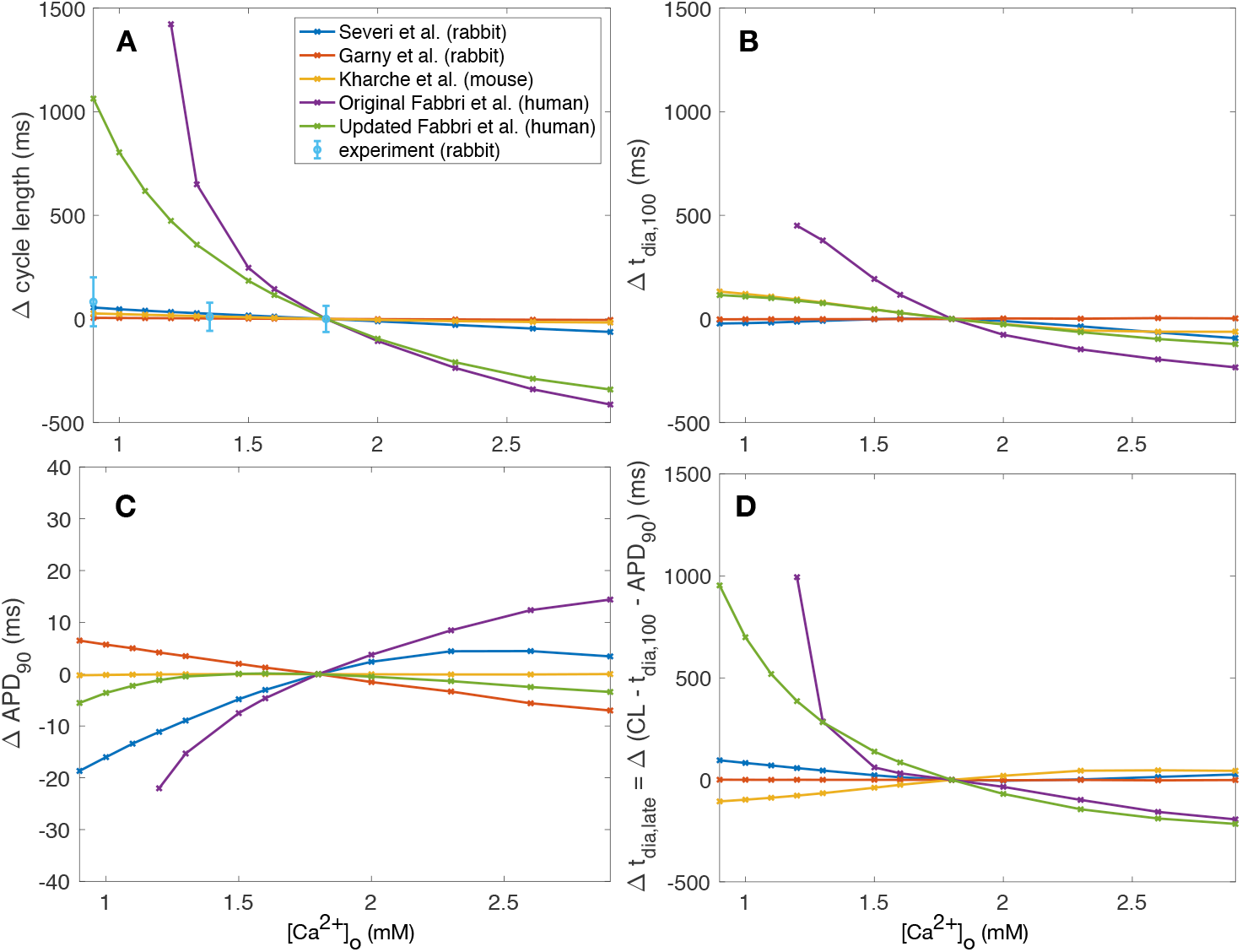
Contribution of different temporal phases of the action potential to the dependence of pacemaking cycle length (A) on extracellular calcium concentration changes (reference: 1.8 mM) in models of different species: first order approximation of diastolic depolarization based on early depolarization (100 ms, B), action potential duration at 90% repolarization (C), diastolic depolarization not covered by first order approximation based on DDR_100_ (D).

We dissected the contribution of the different phases of the pacemaking cycle to the cylce length dependence on [*Ca*^2+^]_*o*_ (Fig. 3A). The effect of early depolarization is quantified by calculating a first order approximation of diastolic depolarization duration based on DDR_100_ and a takeoff potential of −40 mV:

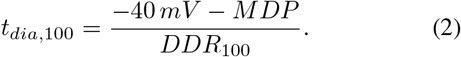

*t_dia,late_* (Fig. 3D) on the other hand quantifies all changes that are captured by neither changes in early depolarization (*t_dia,_*_100_, Fig. 3B) nor by changes in *APD*_90_ (Fig. 3C). While *APD*_90_ changes are responsible for only a minor share of the overall effect, early depolarization becomes slower under hypocalcaemic conditions in the Kharche et al. mouse model and, most pronounced, in the human models. While this effect is counterbalanced by *t_dia,late_* in the mouse model, this is not the case for the human models. Late depolarization becomes more important for [*Ca*^2+^]_*o*_ < 1.5 mM, particularly for the updated human model.

## IV. DISCUSSION

In this paper, we identified crucial inter-species differences in the response of cellular pacemaking in the sinus node to changes of extracellular ion concentrations, as commonly occurring in CKD patients. The bradycardic effect of hypocalcaemia was markedly more pronounced in the human models compared to small animal models. The pronounced effect is still present in the proposed updated version of the Fabbri et al. model of human sinus node cells, which exhibits homeostasis of intracellular ion concentrations across time spans of minutes and thereby puts further physiological constraints on the model parameters compared to the original version without impairing reproduction of experimental AP and CaT features. The similar behavior of the Fabbri et al. model variants and the Severi et al. model for APD_90_, (dV/dt)_*max*_, and overshoot could be due to the fact that the latter served as basis to derive the former model. When interpreting the effect in terms of absolute [*Ca*^2+^]_*o*_, one should keep in mind the difference between in vitro and in vivo reference concentrations (1.8–2.0 mM vs. 1.0–1.3 mM) [14]. Future studies could investigate the contribution of distinct currents and their complex interplay. Moreover, the analysis of autonomic regulation as well as pace-and-drive capability [15] adds other dimensions to the problem of bradycardic SCD.

We conclude that the relevance of mouse and rabbit experiments to investigate the effect of electrolyte-induced changes of pacemaking appears questionable and human experiments are needed to investigate the potential mechanism of hypocalcaemia-induced bradycardic SCD in CKD patients.

